# The U.S. faculty job market survives the SARS-CoV-2 global pandemic

**DOI:** 10.1101/2022.05.27.493714

**Authors:** Ariangela J Kozik, Ada Hagan, Nafisa M Jadavji, Christopher T Smith, Amanda Haage

## Abstract

**Purpose:** This paper aims to identify the extent to which the COVID-19 pandemic disrupted the academic job market and the ways in which faculty job applicants altered their applications in response to a changing academia.

**Design/methodology/approach:** The data presented here is the portion relevant to COVID-19 collected in a survey of faculty job applicants at the end of the 2019-2020 job cycle in North America (spring 2020). An additional “mid-pandemic” survey was used in fall 2020 for applicants participating in the following job search cycle to inquire about how they were adapting their application materials. A portion of data from the 2020-2022 job cycle surveys was used to represent the “late-pandemic”. Job posting data from the Higher Education Recruitment Consortium (HERC) is also used to study job availability.

**Findings:** Examination of faculty job postings from 2018 through 2022 found that while they decreased in 2020, the market recovered in 2021 and beyond. While the market recovered, approximately 10% of the faculty job offers reported by 2019–20 survey respondents were rescinded. Respondents also reported altering their application documents in response to the pandemic as well as delaying or even abandoning their faculty job search.

**Originality:** This paper provides a longitudinal perspective with quantitative data on how the academic job market changed through the major events of the COVID-19 pandemic in North America, a subject of intense discussion and stress, particularly amongst early career researchers.

## Introduction

In the life cycle of academic research, the transition from trainee to independence is generally marked by securing a tenure-track faculty position. The ‘faculty job market’ is a collective term that refers to the pool of highly-qualified academics competing for a limited number of open positions during any given year. In North America, the process generally begins with the announcement of open positions in the summer with application deadlines in the fall and a series of interviews culminating in offers sometime in the late spring the following year. The application process also generally requires job seekers to submit a portfolio comprised of several documents that describe their accomplishments, track record of funding, research and teaching plans, letters of support from recommenders, and sometimes commitments to diversity and inclusion. These documents are tailored for each application, taking into account the stated priorities of the departments and institutions seeking to hire (process reviewed in more depth (Fernandes *et al*., 2020))

In the 2018-2019 cycle, our group sought to determine what it takes to get a faculty job in terms of quantifiable measures of scholarly metrics, application process, funding history, teaching experience, and the interplay between these variables, largely in the life sciences and largely at research intensive universities. Our data revealed that obtaining certain thresholds of scholarly metrics, like six first author publications, or thresholds of the application process, like applying to at least 15 openings, were significant gateways to outcome success, but no clearly defined path exists from trainee to independent faculty position. Overall, integrating quantitative and qualitative data from this foundational survey allowed us to conclude that applicants perceive the market to be elusive, time-consuming, stressful, and limited in feedback (Fernandes *et al*., 2020).

In January 2020, in the midst of the 2019-2020 job cycle, the United States recorded their first case of COVID-19, a disease caused by the virus SARS-CoV-2 (Figure 1). Most states in the US enacted stay-at-home orders by spring 2020, with regionally variable mitigation measures in place throughout 2020 and 2021 (Baker *et al*., 2020). The COVID-19 pandemic impacted nearly every sector of society, including higher education. The University of Washington was the first institution to close its campus on March 7, 2020. Harvard University followed days later, evacuating student residences (Hess, 2020). The spring and summer of 2020 saw the closures of institutions bankrupted after mandatory stay-at-home orders (Cartolano, 2020; Nast, 2020) and remaining institutions enacted austerity measures in response to the unprecedented financial conditions, many of which targeted hiring. Unofficial reports cited at least 400 instances of frozen or canceled faculty searches in the 2019-2020 cycle and beyond, as well as other impacts on hiring (“The Professor Is In. - Posts | Facebook”, n.d.), which many perceived as further destabilizing the already precarious academic market (“Scores of colleges announce faculty hiring freezes in response to coronavirus”, n.d.).

**Figure 1.**
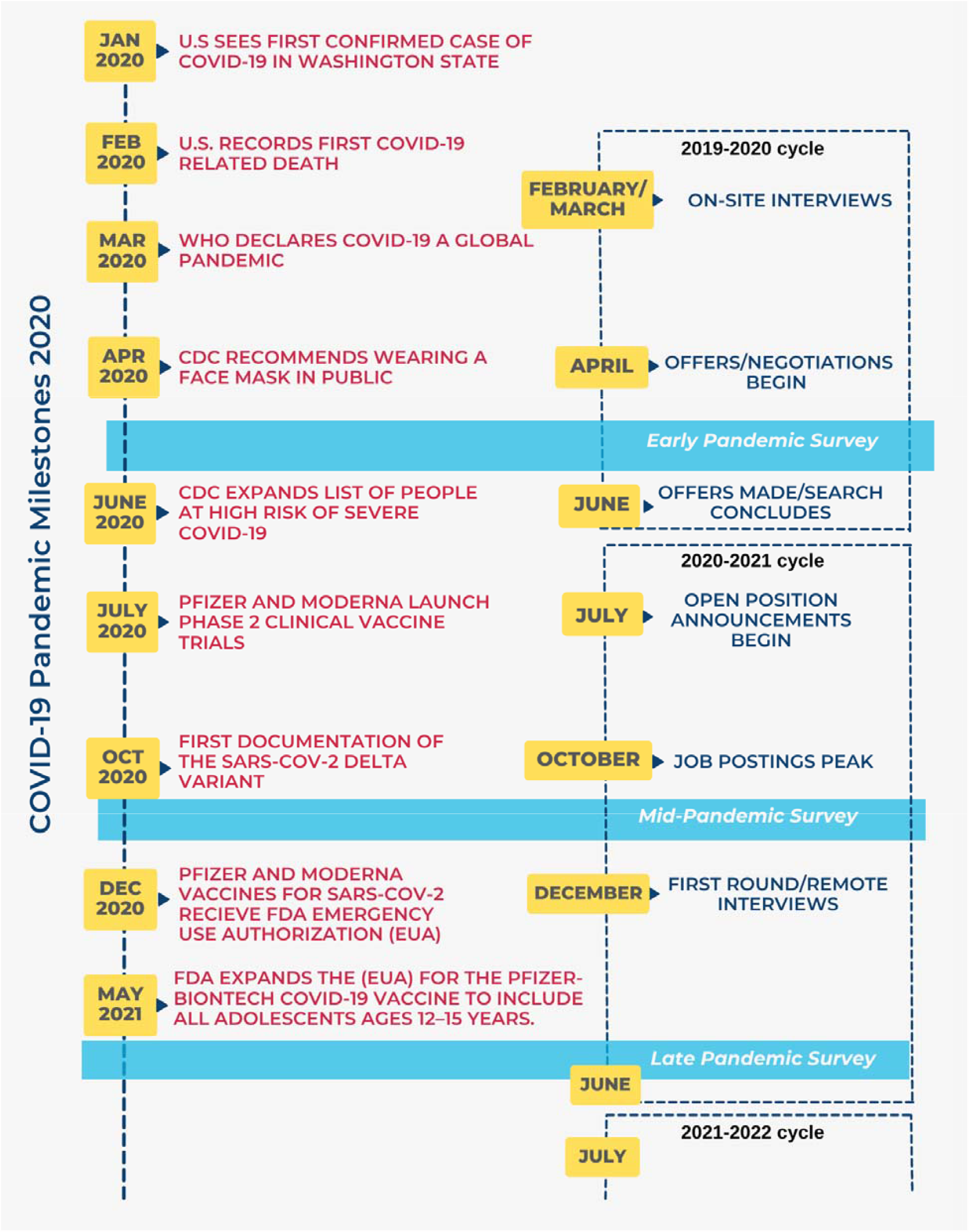
Summary of the faculty job market with COVID-19 pandemic milestones. The COVID-19 pandemic impacted the faculty job cycle with campus closures and stay-at-home orders occurring during the on-site interview period. We administered survey questions (presented/discussed below) at three points, May 2020, November 2020, and May 2022.

As of March 2023, official tallies reported over 675 million cases and 6.8 million deaths worldwide from COVID-19 (“COVID-19 Map - Johns Hopkins Coronavirus Resource Center”, n.d.). The US administration announced an official end to the national and public health emergency as of May 2023, and data is beginning to emerge on how the COVID-19 pandemic impacted early career academics across numerous fields. Recently reviewed by Lokhtina et al., the four main areas affected by COVID-19 include 1) research activity, 2) researcher development, 3) career prospects, and 4) well-being (Lokhtina *et al*., 2022). Of note, a vast majority of studies included in this review, and others (Herman *et al*., 2021) focus on self-reported perceptions in these listed areas. This is particularly prominent in the discussion of “career prospects’’. For example, in a National Postdoc Survey conducted by the University of Chicago, a majority of respondents rated the job market as poor or fair, which represented significant decreases compared to pre-pandemic data, with a possible uptick in decisions to change career trajectory among the survey population (Morin *et al*., 2022). In a similar, widely cited Nature survey, approximately two thirds of postdocs reported that the pandemic negatively affected their career prospects, some citing “suspected” canceled job opportunities as the reason for this negative effect (Woolston, 2020). In several specific fields, like animal behavior and welfare (Sarah *et al*., 2021), clinical sciences (Doyle *et al*., 2021), marine science (Schadeberg *et al*., 2022), and developmental origins of health and disease (Bansal *et al*., 2022), surveys report the increased concerns of early career academics for their future prospects in and outside of academia.

While some of these areas, like well-being, are confined to individual perspective, others can be reflected in quantifiable measures. For example, a sharp decrease in the number of pre-printed manuscripts shortly after the onset of the COVID-19 pandemic compared to expected values for gender and region on the life sciences open access server, BioRxiv, is a proxy measure for research activity or productivity (Abramo *et al*., 2021). With our previous quantitative approach in mind, we sought to answer several research questions overall aimed at quantifying the degree to which the COVID-19 pandemic disrupted the academic job market.To our knowledge at the time of writing, this is the only report to quantify rescinded job offers and combine applicant perception/outcome data with job availability.

**RQ1: How did the pandemic affect the immediate 2019 - 2020 faculty job cycle?**

**RQ2: How did the pandemic affect the availability of faculty jobs?**

**RQ3: How are applicants navigating the faculty job market differently in the wake of the pandemic?**

## Methods

### Data Collection

We designed a survey (the early pandemic survey) to collect self-reported demographics and academic metrics for assistant professor applicants during the 2019-2020 academic job search cycle. The survey was open from May 6, 2020 to September 3, 2020, and respondents (n=791) were not required to answer all questions. After collecting and performing an initial analysis of the early pandemic survey, we designed an additional mid pandemic survey. The mid pandemic survey was open from November 12, 2020 to January 12, 2021 and targeted potential 2020–21 assistant professor applicants (n=78). Finally, a late pandemic survey collected assistant professor applicant outcomes during the 2020-2021 and 2021-2022 job cycles, it was available from May to September 2022. Variables of interest for the early and late pandemic surveys included faculty application outcomes such as interviews, offers, and rescinded offers as well as the corresponding institutions; applicant offer responses; applicant response to the COVID-19 pandemic; as well as the applicant demographics of gender, race, research category, position, and first-generation PhD status. Variables of interest for the mid pandemic survey included plans to apply for assistant professorships in the 2020-21 academic hiring cycle, current position, research category, desired institution of employment (either primarily undergraduate institution [PUI] or research intensive [RI]), extent of changes to research and teaching statements, and the impact of the COVID-19 pandemic on their plans for applications, interviews, and offer responses.

All surveys were distributed by several postdoctoral association mailing lists in North America, Europe and Asia as well as on various social media platforms including the Future PI Slack group, Twitter, and Facebook. All surveys were approved by the University of North Dakota’s IRB office under project number IRB0003954.

Job advertisement data from January 2018 to December 2022 were generously provided by the Higher Education Research Consortium. Variables of interest included the institution name, institution location, hiring position title, position category, the number of positions available, position description, and the dates of advertisement availability. Aggregated data and survey questions are available in the GitHub repository: https://github.com/Faculty-Job-Market-Collab/Kozik_COVID-19_SGPE_2023.

### Data Categorization & Analysis

Where institutions were named (early and late pandemic surveys and job advertisement data), the institution names were cleaned manually and joined with the 2018 Carnegie classification data (“Carnegie Classifications | Downloads”, n.d.). Using these data, we classified educational institutions based on the National Science Foundation definition for PUIs as colleges and universities that have awarded 20 or fewer PhD/DSci degrees during the previous academic year. An institution with more than 20 PhD/DSci degrees awarded during the previous academic year was classified as an RI institution. Both the Carnegie classification and job advertisement location data were used to identify institution country and region within the USA (Table S1), where appropriate.

Survey respondents who indicated that they were either non-binary, transgender, or that their gender was not listed were grouped as gender minorities (Trans/GNC). Respondents were allowed to select multiple race/ethnicity categories, which were grouped into two categories: persons historically excluded due to ethnicity or race (PEER) and non-PEER. PEER respondents were those who self-identified with one or more of the following identities: Oceania, Not Listed, North American Indigenous, Caribbean Islander, North American Hispanic/Latinx, and Black/African American/African. The following research fields were grouped together under “Mathematics & Engineering Sciences”: Mathematical & Physical Sciences, Engineering, and Computer & Information Sciences. Humanities and Social, Behavior, & Economic Sciences were grouped together under “Social & Behavioral Sciences”. A positive response to changes in research or teaching statements included either “yes, significant changes” and “somewhat” or “strongly agree”, “agree”, and “somewhat agree”, as appropriate.

Job advertisements were considered tenure-track positions if “tenure track” was referenced in either the position title or description. Tenure-track positions were then categorized as early career research or assistant professor positions according to the presence of “assistant” in either the position title or description. Advertisements for temporary positions contained the terms “non tenure”, “fixed-term”, “adjunct”, “lecturer”, or “instructor” in either the position title or description.

Data were manipulated and visualized using R statistical software (version 4.3.1) and relevant packages. All code used for data analysis and visualization are available in the GitHub repository: https://github.com/Faculty-Job-Market-Collab/Kozik_COVID-19_SGPE_2023.

### Statistical Analysis

Between group differences were tested using the X^2^, Fisher’s exact, or one-way ANOVA tests for statistical analysis using R. Wherever statistical analyses were used, the tests and p-values are reported in the corresponding figure legend. The level of significance was set to p < 0.05.

## Results

### RQ1: How did the pandemic affect the immediate 2019 - 2020 faculty job cycle?

The COVID-19 pandemic collided with higher education in March 2020 (Figure 1). In terms of the 2019 - 2020 North American tenure-track faculty job market, this is fairly late in the applicant interview process. In-person interviews were likely nearly completed and verbal to written offers were beginning to be negotiated. Our first research questions asked what abrupt changes occurred to this process as the uncertainty of a global pandemic unfolded. To answer this we asked respondents from a 2019 - 2020 iteration of a larger on-going survey of faculty job search outcomes whether the pandemic affected their search that cycle (early pandemic survey, see methods).

Given where the cycle was at when the pandemic’s impacts were first felt, we first looked at how many on-site interviews were moved to remote. The on-site interview usually follows a video conference or phone interview that took place weeks or months earlier. During the on-site interview, faculty candidates usually provide critical presentations on their past and future work as well as meet a majority of the department and see the prospective institution and surrounding area, likely for the first time. In the 2019 - 2020 cycle, 13% of on-site interviews were moved to off-site interactions. While an established rate at which this occurs prior to the pandemic is not available in the literature, we have asked this question in subsequent survey cycles. In comparison, 83% of on-site interviews were conducted off-site in the 2020 - 2021 cycle and 35% were conducted off-site in the 2021 - 2022 cycle. This is congruent with the timing of pandemic events as scheduling for on-site interviews for the 2020 - 2021 cycle would have likely occurred prior to or during vaccine rollout, during institution stay-at-home orders. This repeated pattern of a higher level of remote interactions in the 2021 - 2022 cycle likely reflects some continued value of remote and/or flexible work in higher education.

As previously mentioned, it is very likely that offer negotiations were well underway in a majority of faculty searches in March 2020. Early pandemic survey respondents (2019 - 2020 cycle) reported that 10.3% of the offers received were rescinded by the institutions. Again, an established rate of rescinded offers each cycle prior to the pandemic is missing in the literature, but we have maintained this question on our survey iterations. The percent rescinded dropped to 7.9% and 0.7% in the 2020 - 2021 and 2021 - 2022 cycles, respectively. This notes a significant disruption in the faculty job market amid the COVID-19 pandemic, thus we wanted to analyze rescinded offers by applicant and institutional demographics, particularly for the early pandemic 2019 - 2020 cycle. More men (18.3%) reported having an offer rescinded compared to women (14.6%) and gender minorities (7.7%), though the results were not statistically significant (Figure 2A; p = 0.54). We also observed no significant difference in percent offers rescinded by PEER status (Figure 2B; p = 0.65), defined as persons excluded because of their ethnicity or race through the history of racism in the United States (Asai, 2020), research category (Figure 2C; p=0.54), type of institution where the offer was made (Figure 2D; p=0.61), or geographic region where the offer was made (Figure 2E; p=0.16).

**Figure 2.**
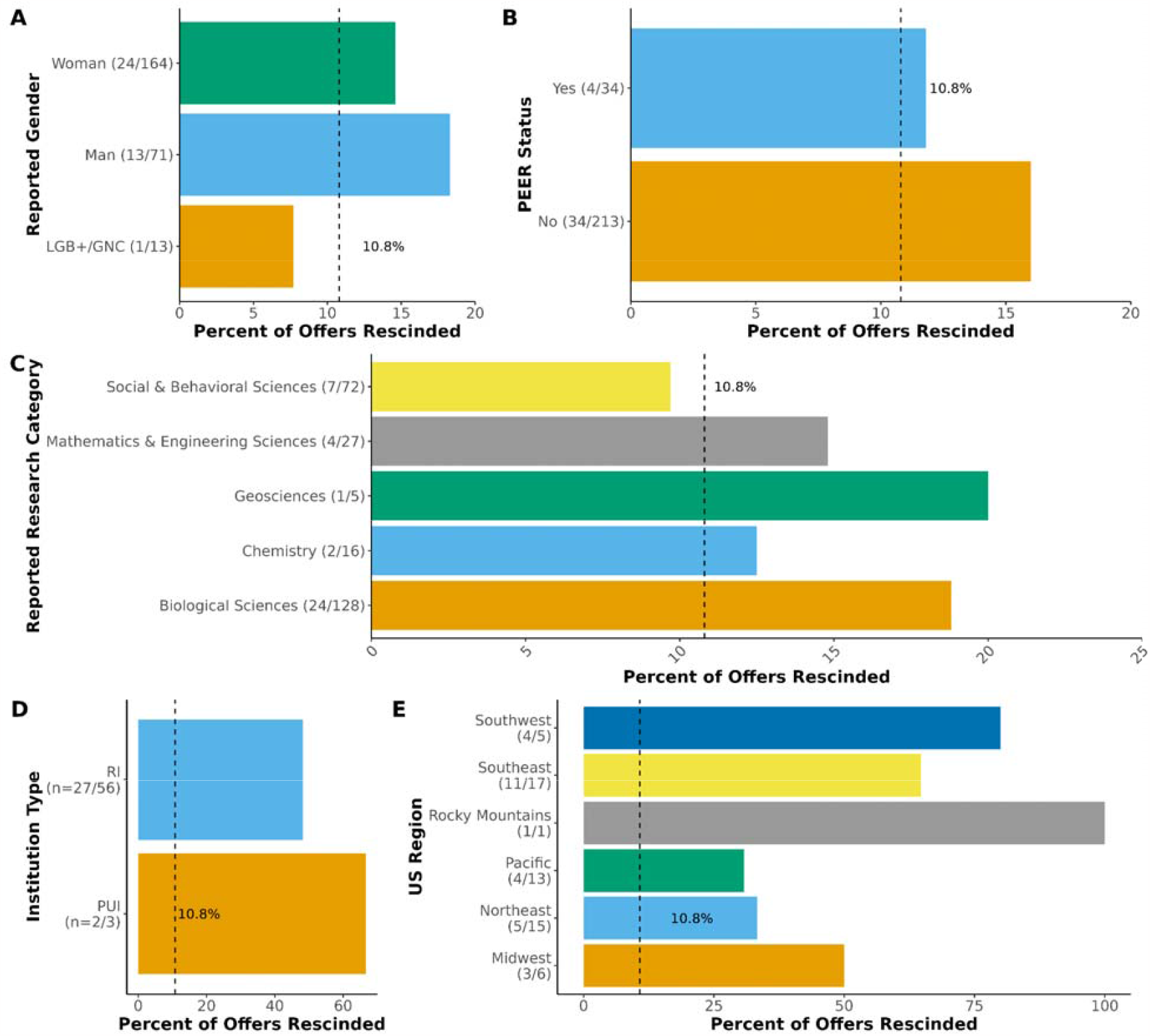
The percent of assistant professor job offers rescinded during the 2019-2020 hiring cycle according to applicant and institutional demographics. Survey respondents who indicated that they received faculty offers were asked to name the institutions that they received offers from and to indicate how many of those offers were rescinded. The percent of offers rescinded were calculated based on the respondent’s: A) Reported gender (p=0.54), B) PEER status (p=0.65), C) Reported research category (p=0.54), D) US institution type (p=0.61), and E) the US region of the institution extending the offer (p=0.16). The vertical, dashed black line indicates the percent of offers rescinded across all responses. Non-responses to the question regarding rescinded offers were omitted from the analysis. In parentheses, the number of offers rescinded/the number of offers. P-values obtained using Fisher’s exact test.

We also asked survey respondents about rejecting faculty job offers for a variety of reasons, including because of “reduced resources directly related to the economic impact of COVID-19” or “other reasons pertaining to the impact of COVID-19”. The overall rate of rejected offers in the 2019 - 2020 cycle was 18.3% with 12% of those being due to COVID-19-related reasons. Again, the rate at which candidates reject faculty job offers overall remains unknown in prior cycles, but in our subsequent surveys the rejection rate was 19.2% and 23.8% in the 2020 - 2021 and 2021 - 2022 cycles, with 9.1% and 2.7% of those being due to COVID-19, respectively. Interestingly, this follows a similar trend downward as rescinded offers. Together this data indicates there were significant immediate disruptions to the North American faculty job market, particularly in offers being rescinded late in the cycle and candidates had to make difficult decisions when accepting offers amidst uncertainty.

### RQ2: How did the pandemic affect the availability of faculty jobs?

While offers rescinded in spring 2020 were certainly devastating to the affected individuals (Langin, 2020), the largest impact was projected as yet to come as some viewed the pandemic as an “extinction event” for academia (https://www.chronicle.com/author/karen-kelsky, 2020). In fall 2020, postdocs reported an increasingly negative perception of the academic job market due to the pandemic and many delayed their search accordingly (Morin *et al*., 2022). To understand if these fears aligned with the data, we analyzed job posting data from 2018 through 2022 collected by the Higher Education Recruitment Consortium (Figure 3). The number of tenure-track positions of all levels were consistently lower in 2020 compared to the average of all other years examined. They started low, lagged throughout the fall, and never quite reached the peaks seen in previous or subsequent years (Figure 3A). A similar trend was seen when we investigated assistant professor tenure-track positions alone (Figure 3B). This lag was previously described by *Science* as the market “tanking”, using data through September 2020 (Langin, 2020), but the 2021 and 2022 posting cycles appear equivalent to, if not higher than, the 2018 and 2019 cycles, suggesting that the number of Assistant Professor and All Tenure-Track positions bounced back after a period of uncertainty (Figures 3A & 3B, Supplementary Figure 1).

**Figure 3.**
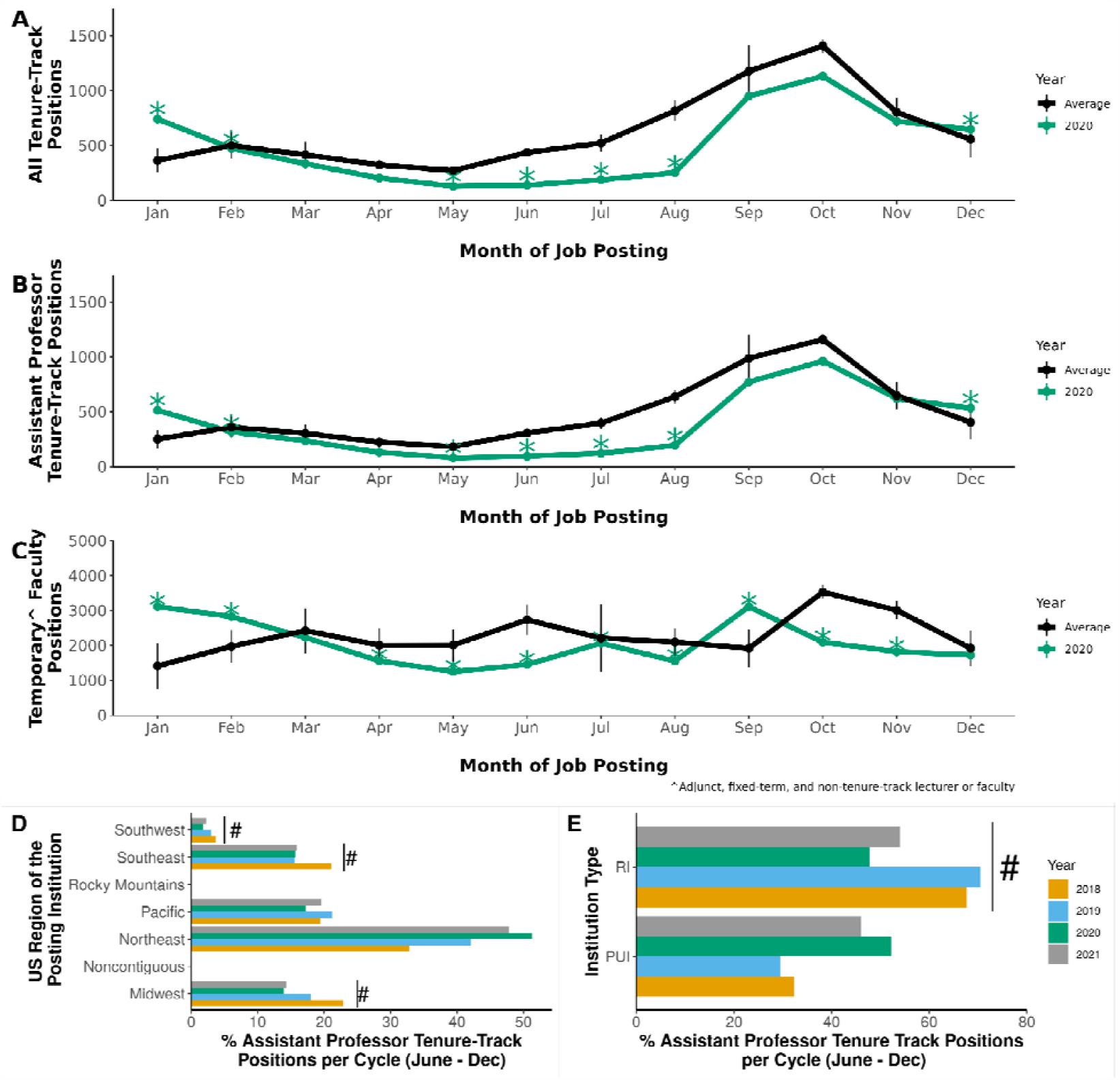
The impact of the COVID-19 pandemic on the academic job market. Job posting data from January 2018 to December 2022 were obtained from the Higher Education Recruitment Consortium (HERC). The years 2018, 2019, 2021, and 2022 were averaged (black) by month and compared to 2020 (green) according to the job type: A) All tenure-track positions, B) assistant professor tenure-track positions, and C) temporary faculty positions, which included adjunct, fixed-term, and non-tenure-track lecturers or faculty (not postdoctoral positions). The percent of positions posted for each year (June to December) from 2018 to 2021 were plotted according to the D) US region and E) institution type (RI or PUI). P-values obtained using (A-C) Pearson’s X^2^ test with Bonferroni correction and (D-E) one-way ANOVAs with a Tukey multiple comparison of means. p<0.001 = ^*^; p<0.05 = #. Vertical lines (A-C) indicate 1 standard deviation.

COVID-19 also had a significant impact on the number of temporary faculty positions. Perhaps unsurprisingly, the number of these positions and their trends throughout the calendar year are generally more erratic than tenure-track positions (Supplementary Figure 1). Though the number of temporary positions started off high in 2020, they quickly dropped once the pandemic was evident in the USA, to unprecedentedly low levels (Figure 3C). The number of temporary faculty positions posted between April and August 2020 is the absolute lowest seen compared to 2018, 2019, 2021, and 2022. Interestingly, 2022 may be the highest year in recent history for the number of temporary faculty positions posted (Supplementary Figure 1). Responses to the interruption in the market also differed by institution type and region, pointing to a significant level of year-by-year variability in what type and where jobs are available. (Figures 3D & 3E).This inherent variability in the faculty market likely contributes to the feelings of unease and opaqueness by participants (Fernandes *et al*., 2020). Despite this, data on job postings and the downward trend in rescinded and rejected offers overall demonstrates that even though COVID-19 likely contributed to a significant disruption in job availability in late 2019 and throughout 2020, this impact was acute and the market is largely similar to before.

### RQ3: How are applicants navigating the faculty job market differently in the wake of the pandemic?

While the number of faculty job postings appears to be back to pre-pandemic levels, many speculate that what a faculty job is, the roles and responsibilities, and how they are carried out, is likely forever impacted by the practices adopted to accommodate pandemic safety measures. Teaching an online modality to accompany a traditional in-person section of a class (“Growth in online education. Are providers ready? | McKinsey”, n.d.; Morris *et al*., 2020), or teaching in a hybrid manner may now be an expectation (“Staying Relevant”, n.d.), despite pedagogical criticisms (Gillis, n.d.). At a minimum, flexibility in course policies and practices is the new norm (https://www.chronicle.com/author/beckie-supiano, 2023). Research expectations continue to climb (Strawser, n.d.), with highly variable institution responses to disruptions, such as tenure extensions or changes in policy (Butler, n.d.). In addition, institutions continue to explore how remotely working staff or faculty fit into their business models (https://www.chronicle.com/author/lindsay-ellis, 2021; Lederman, n.d.; Ph.D, n.d.). Due to these cultural shifts, we wondered how applicants may change their approach to the market, particularly in the focus of their prepared documents outlining their teaching and research plans, and their commitments to achieving a faculty job. To answer this we compared two of our larger survey iterations, the 2019 - 2020 cycle with the “early pandemic respondents” and the 2020 - 2022 with the “late pandemic respondents” with an additional mini climate survey deployed in fall 2020 with “mid pandemic respondents”, aimed at those in the midst of applying during arguably the height of pandemic affects (see methods).

First, we asked early (Figure 4A) and late (Figure 4C) pandemic respondents if they “altered their research statement to focus on remote or computational research”. Unsurprisingly this rate is fairly low (5%) in early pandemic respondents (Figure 4A) as pandemic impacts rolled out across the nation late in the cycle, likely after applicants prepared and submitted most of their written material. The frequency of altering to a more remote or computational work increased in the late pandemic respondents to an average of 14.6% (Figure 4C). This is reproduced in the mid pandemic respondents (Figure 4B) who were asked if they altered their research statements to “be more remote-friendly”, which an average of 22.2% said they had. Additionally, mid pandemic respondents also indicated that they “incorporated pandemic-related topics” into their research statements at relatively the same rate (Figure 4B).

**Figure 4.**
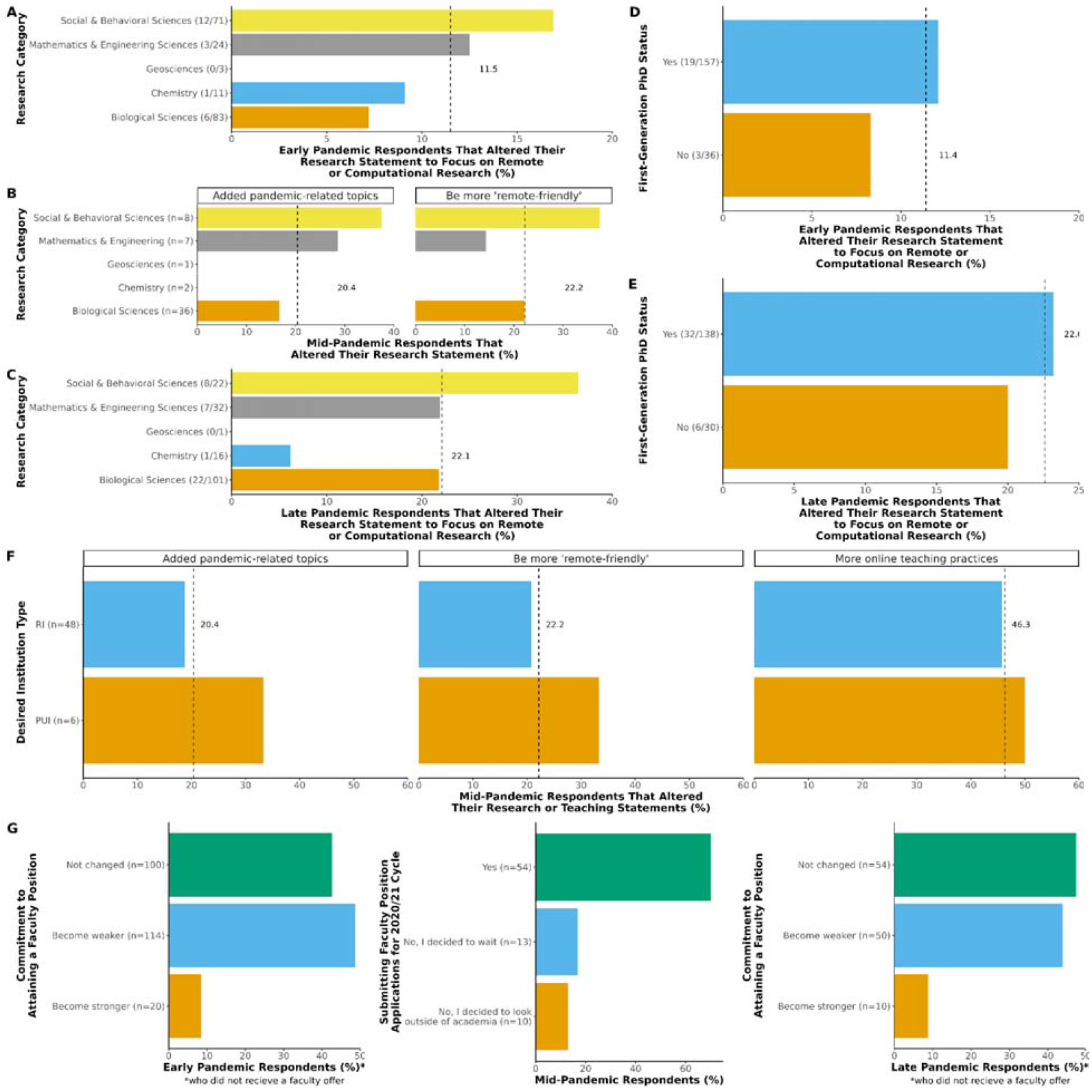
A comparison of early, mid, and late pandemic attitudes on search strategy and academia. The percent of respondents in each research category from A) the early pandemic survey that altered their research statements to focus on remote or computational research (p=0.43), B) the mid pandemic survey that altered their research statements to (*right panel*) include pandemic-related areas (e.g., coronavirus or COVID-19) when they would not have otherwise done so (p=0.59) and/or (*left panel*) to be more “remote-friendly” (p=0.81), and C) the late pandemic survey that altered their research statements to focus on remote or computational research (p=X). The percent of D) early pandemic and E) late pandemic respondents that altered their research statement to focus on remote or computational research based on their first-generation PhD status (p=0.43 and p=X, respectively). F) The percent of mid-pandemic respondents interested in either RIs or PUIs that altered their research statements to (*left panel*) add pandemic-related research (p=0.59) or (*center panel*) be more remote friendly (p=0.6) and/or (*right panel*) those who altered their teaching statements to include more online practices (p=1). G) The responses (%) of early pandemic respondents (*left panel*) who did not receive faculty offers when asked about their commitment to attaining a faculty position; mid pandemic respondents (*center panel*) when asked about their plans to submit faculty position applications during the 2020-2021 hiring cycle (p<0.001); and late pandemic respondents (*right panel*) who did not receive faculty offers when asked about their commitment to attaining a faculty position. n = the number of respondents. The vertical, dashed line (black) indicates the percent for all groups pooled together. P-values obtained from (A-F) Fisher’s exact test and (G) the X^2^ test for given probabilities

We also looked for trends in who might be more likely to alter their research statements across field (Figure 4A-C), institution type to which applicants were primarily applying (Figure 4F), or possibly experience in higher education, which we approximated by comparing responses of first-generation applicants (Figure 4D & E). Though not always statistically significant, there was an apparent trend in the fields of respondents that adapted their work to be more remote, with geosciences and chemistry consistently being lower than other fields (Figure 4A-C), possibly due to the nature of research in these fields and their inherent flexibility. A trend also appeared when comparing institution type, where mid pandemic applicants to primarily undergraduate institutions (PUIs) appeared to “incorporate pandemic-related topics” (33.3%) and alter their research statements to “be more remote-friendly” (33.3%) at higher rates compared to applicants to R1 institutions (∼20% for both measures) (Figure 4F). This may reflect the purpose of research and funding sources at these types of institutions. Research at PUIs generally serves mostly to introduce undergraduates to the process of research and thus has less federal funding. RI, or research intensive institutions more highly value consistent research output as a large portion of their operational model requires high amounts of external grant funding. Lastly, faculty job applicants who were the first in their family to achieve a PhD had a significantly higher rate of altering their research statements both in the early and late in the pandemic surveys. Specifically in the late pandemic survey 25.6% of first-generation PhD holders versus 14.3% of their non-first generation peers altered their research statement to include more computational or remote work (Figure 4D; p=7.13E-05). It is interesting to speculate whether this may reflect an expectation of those less experienced for significant change, or generational knowledge of stable, conservative institutions.

Though altering the research statement to be more readily remote was done at a relatively low rate across any time through the pandemic (∼20%), mid pandemic applicants in the fall of 2020 appeared to more readily change their teaching philosophy statements to include “more online teaching practices” with an average of 46.3% positive responses. (Figure 4F). This was also comparable between institution types with 46% of RI and 50% of PUI applicants including an aspect of online teaching in their materials. Anecdotally this appears consistent with the discussion of higher education throughout the pandemic, as it largely focused on the emergency transition to remote teaching and the after effects. Finding similar opinion pieces on the flexibility in research demands was difficult.

While studying how applicants changed their written materials tells us some of what applicants are anticipating in terms of changes to the roles and responsibilities of being a faculty member, it does not give us a picture of the outlook applicants have on the market overall. Previous work has indicated that applicants view the market as stressful and opaque (Fernandes *et al*., 2020), with decreasing outlook due to pandemic effects (Morin *et al*., 2022; Woolston, 2020). We asked early pandemic (2019 - 2020 cycle) and late-pandemic (2020 - 2021 cycle) applicants about their “commitment to attaining a faculty position” and mid pandemic (fall 2020) applicants about their potential applications in the cycle to measure this. Our mid pandemic survey specifically asked respondents: “Are you currently applying for faculty jobs in the United States or Canada with an anticipated start date in 2021 or 2022?”. The difference in response rates to this question were statistically significant (X^2^=44.97, df=2, p<0.001). Approximately 69% of respondents were applying during the 2020-21 job cycle while 16.7% decided to wait and 12.8% decided to change career paths and look for employment opportunities outside academia. These data are consistent with the 11% of respondents to the Morin, et al study who reported delaying their job search due to the pandemic (Morin *et al*., 2022). Interestingly, the early-pandemic and late-pandemic responses to “After participating in the academic job market this year, my commitment to an academic career has:” was not significantly different with ∼40-50% responding that their commitment hadn’t changed, another ∼40-50% saying that it had become weaker, with the last small percent responding that their commitment was now stronger. Together this data indicates that the largest change applicants are making to navigating the faculty job market is incorporating online teaching into their written materials, but overall it appears that business as usual has returned to the market.

## Discussion

Our ongoing efforts to curate data pertaining to the faculty job market provided a unique opportunity to observe the real-time impact of the SARS-CoV-2 pandemic and the accompanying global uncertainty and disruption. This analysis provides critical information to efforts to understand applicant decision making in an increasingly competitive market among sustained atypical conditions.

We quantified, for what we believe to be the first time, the percent of rescinded faculty offers in the 2019 - 2020 application cycle at ∼10%, as the pandemic first influenced higher education. Despite this negative outcome for some, the rate at which respondents retained their commitment to the faculty job market at the end of the cycle remained similar to later years (∼45%). We also demonstrated a reduced number of faculty job postings throughout all of 2020 by 20 to 30%, which is supported by other analyses (Langin, 2020; Olena, n.d.; Woolston, 2020). However, our analysis of the ‘post-pandemic’ period is among the first to suggest that the market has indeed found a new equilibrium, with a more consistent number of postings in 2021 and 2022. In addition, ∼16% of respondents chose to not apply during the 2020 - 2021 cycle and wait the uncertainty out. The number of job postings in 2021 and 2022, while going back up to pre-pandemic levels, did not reach 20 to 30% higher to make up the positions lost in 2020. These numbers, though possibly deceptively small when examined in isolation, altogether paint a picture of significant loss and delay in the faculty job market. For example, if 45% of the 10% who had their offers rescinded in spring 2020 then applied again in fall 2020, this places increased competitive pressure on an up to 30% reduced pool of positions. Estimating this at 45% is likely generous, as applicants who made it to the offer stage are highly successful and may have an increased sense of commitment given their successful progression to the late stage of a search in a previous cycle. The increase in the applicants for Fall 2020 then may be somewhat alleviated by the 16% who chose not to apply that cycle, but not completely, and this may just pass the competitive pressure on to future cycles. Conversely, another likely impact of the disruption was the exodus of existing faculty and postdocs from academic science altogether during the pandemic period. This could have eased competition slightly as the applicant pool decreased. Our data also demonstrated the year-to-year variability in faculty job opportunities across region and institution type such that any disruption causes a potentially devastating loss of unique positions. Though an area to be explored more in future work, it is clear that applicants have highly personalized reasons for rejecting offers, and without the right position, some talented applicants who encountered this onerous three year faculty job search landscape were undoubtedly lost. The faculty job market is already considered by many to be unduly competitive, stressful, lacking in tangible feedback, and taxing on applicants’ resources. This increasingly unclear path to the typical outcome (a tenure-track position) many are trained for throughout their PhD or postdoctoral positions is antagonistic to a sustainable future for academia. There have already been reports of excessive difficulty in recruiting postdocs, with many new PhD graduates opting to directly transition to other sectors (Wosen, 2023). Pandemic-initiated conversations across social media and in major organizations like the NIH about the structural problems in the academic ecosystem have also likely discouraged many from participating at all. Therefore, the impact of this seemingly acute temporary disruption in offers, position numbers, and applicant commitments will be felt in faculty recruitment for years to come.

An unanticipated result of our study was the relatively low percent of respondents who adapted their application materials. The highest response to altering application materials was ∼45% of applicants in the fall 2020 included “more online teaching practices” in their teaching philosophy statements. Though this is relatively high compared to the amount who adapted their research plan, this is still relatively low compared to the number of faculty who had to teach online during at least part of 2020, which we estimate to be upwards of 90%. Interestingly, even as the pandemic progressed and society saw the full pre and post vaccine impacts, only 14.6% of applicants in the 2021 - 2022 cycle adapted their research statement to include possible remote or computational work. This seems at odds with how universities have actually returned to function over the last two years. Many opinion pieces point to a growing trend in remote work across many sectors, including higher education. Online education was already growing pre-pandemic and shows no signs of slowing down. It is both a welcomed improvement in accessibility and also a necessary revenue source for institutions of all types. In addition, the day to day life of faculty and staff remains a hybrid, bouncing back and forth between in-person and online commitments. While there are certainly groups, particularly across the United States, calling for a complete “return to normal” it remains unlikely that higher education will ever look like it did in 2019 -- and perhaps it shouldn’t.

Structural inequities in the United States academic sector have long had a negative impact on faculty recruitment, retention, and promotion, particularly for women and gender minorities, those from historically underrepresented backgrounds, and those with caregiving responsibilities. The pandemic has exacerbated these inequities and accelerated the subsequent consequences–the attrition of talented individuals from a system that wasn’t designed to support them. After the significant decrease in faculty job opportunities in 2020, higher education was also impacted by the ‘Great Resignation’: a pattern of resignations in late 2021 (“Higher Education’s Role in the Era of the Great Resignation”, 2021; “Job Openings and Labor Turnover Summary - 2022 M02 Results”, n.d.; N *et al*., 2021). Attributed to widespread dissatisfaction with wages, career trajectory, and lack of social support from employers, the Great Resignation doesn’t appear to have changed the inequitable policies, practices and norms that hamper the ability of many academics to conduct research, publish, and successfully compete for funding. For example, a lack of affordable and reliable childcare and education options coupled with partial vaccine eligibility for children and inconsistent mitigation strategies at daycares and schools all force caregivers to make difficult choices that pit career advancement against safety. Insufficient institutional support for research/teaching loads, untenable supply chain delays, threat and burden of a potentially debilitating illness, repeated periods of isolation, and increasingly limited time to meet inflexible expectations all reinforce these disadvantages. Ironically, those most vulnerable to these systemic holes in support are those most needed to diversify the academy. Swift and strategic investments are required to address these pressures and mitigate lasting damage to such participants in the academic job market. The lasting impact of the pandemic-induced disruption has provided academia with a unique opportunity to evaluate and change the system for the better. It is likely necessary for applicants to be competitive in a condensed pool to remain ahead of these value shifts in higher education.

### Study Limitations

Our study does have several limitations. First, the surveys were shared using social media, which may have resulted in a sampling bias. However, our surveys were also disseminated through official postdoctoral association/organizational listservs to help mitigate potential bias. Furthermore, respondents did not answer all the questions and/or complete their responses, causing potential classification errors. For example, PUI/RI institutions were classified based on the list of universities provided by the respondent, and institution names were frequently abbreviated. Anonymous responses mean that we were not able to follow up with respondents or get paired responses for later surveys, thus each survey represents a distinct pool. In addition, there are slight variations in question wording between themes compared, particularly in reference to Figure 4 across the early-pandemic, mid-pandemic, and late-pandemic surveys. For the early and late pandemic surveys, because these questions are part of a larger survey, question wording is consistent, as well as response rate. The mid pandemic survey was deployed separately from this structure and thus suffers from slightly different wording, as well as decreased response rate. In order to make interpretation as clear as possible, the direct wording on each survey is included in the graphs for Figure 4 as well as in the repository link provided in the methods. Despite these slight variations we believe the thematic conclusions drawn above align with results from other studies. The survey was aimed at and largely distributed to North American faculty job applicants. This limits the application of conclusions drawn to that of the North American, largely United States, faculty job market. While similar dynamics may apply to other markets, in the authors’ opinion this is a necessary limitation as training, names of metrics and materials, and the processes applicants go through vary a surprising amount in other regional markets. Lastly, these questions were part of a larger, novel, exploratory study into the faculty job market. We acknowledge that these limitations make generalizations difficult, but the authors assert that our findings provide important data and insights that are otherwise missing from the pandemic record. It is our hope that this work can form the basis for larger, well-funded future studies on the faculty job market.

## Conclusions

In conclusion we have determined that the faculty job market in the United States has largely survived the COVID-19 pandemic, thereby avoiding the worst case scenario forecasts. However, will applicants continue to survive the increasingly difficult recruitment landscape? In an already hypercompetitive environment, applicants have had offers rescinded, potentially adding years to their search for a stable position. Other applicants delayed their search only to be met with lost positions, never to resurface. This points to an exacerbation of the already poor situation of the faculty job search, leaving many to make difficult career decisions. With the rise of the “Great Resignation” due to a culture of wage dissatisfaction, perceived lack of support by peers and administrators, and funding schemes incapable of keeping up with inflation, it must be asked if the endless pool of applicants will continue and who the system in its current form is selecting for. Future work is necessary to continue examining these long term impacts and push for much needed reform.

## Supplementary

**Table S1.** United States by region.

| US Region | States |
| --- | --- |
| Midwest | Illinois, Indiana, Iowa, Kansas, Michigan, Minnesota, Missouri, Nebraska, North Dakota, Ohio, South Dakota, Wisconsin |
| Noncontiguous | Alaska, Hawaii |
| Northeast | Connecticut, Maine, Massachusetts, New Hampshire, New Jersey, New York, Pennsylvania, Rhode Island, Vermont |
| Pacific | California, Oregon, Washington |
| Rocky Mountains | Colorado, Idaho, Montana, Nevada, Utah, Wyoming |
| Southeast | Alabama, Arkansas, Delaware, Florida, Georgia, Kentucky, Louisiana, Maryland, Mississippi, North Carolina, South Carolina, Tennessee, Virginia, Washington DC, West Virginia |
| Southwest | Arizona, New Mexico, Oklahoma, Texas |

**Figure S1.**
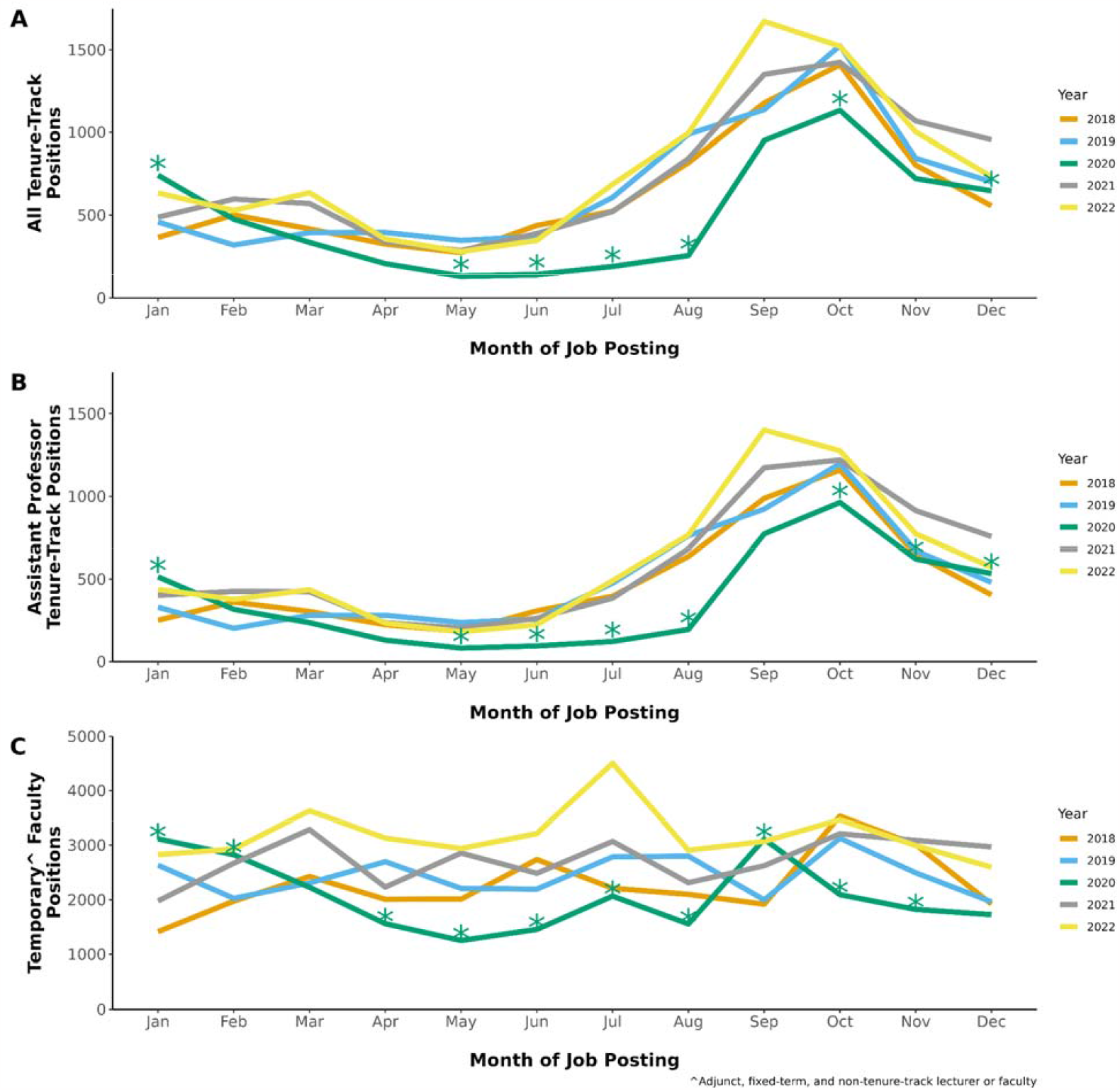
The academic job market by year. Job posting data from January 2018 to December 2022 were obtained from the Higher Education Recruitment Consortium (HERC). The posts for each year were plotted according to the month posted and the job type: A) All tenure-track positions, B) assistant professor tenure-track positions, and C) temporary faculty positions, which included adjunct, fixed-term, and non-tenure-track lecturers or faculty (not postdoctoral positions). P-values obtained using Pearson’s X^2^ test with Bonferroni correction. p<0.001 = ^*^.

## Notes

### Competing Interest Statement

The authors have declared no competing interest.

### Summary of Updates

This version of the manuscript includes a greater discussion of the findings and job posting data from 2022.

